# Asymmetric outcome of community coalescence of seed and soil microbiota during early seedling growth

**DOI:** 10.1101/2020.11.19.390344

**Authors:** Aude Rochefort, Marie Simonin, Coralie Marais, Anne-Yvonne Guillerm-Erckelboudt, Matthieu Barret, Alain Sarniguet

**Affiliations:** INRAE, Agrocampus-Ouest, Université de Rennes 1, UMR 1349 IGEPP, Le Rheu, France; IRHS-UMR1345, Université d’Angers, INRAE, Institut Agro, SFR 4207 QuaSaV, 49071, Beaucouzé, France

**Keywords:** microbiota, coalescence, seed, root, stem, soil, *Brassica napus*

## Abstract

Seed microbial community constitutes a primary inoculum for plant microbiota assembly. Still, the persistence of seed microbiota when seeds encounter soil during plant emergence and early growth is barely documented. Here, we characterized the interchange event or coalescence of seed and soil microbiota and how it structured seedling bacterial and fungal communities. We performed eight contrasted coalescence events to identify drivers influencing seedling microbiota assembly: four seed lots of two *Brassica napus* genotypes were sown in two soils of contrasted diversity. We found that seedling root and stem microbiota were influenced by soil diversity but not by initial seed microbiota composition. A strong selection on the two-source communities occurred during microbiota assembly, with only 8-32% of soil taxa and 0.8-1.4% of seed-borne taxa colonizing seedlings. The recruitment of seedling microbiota came mainly from soil (35-72% of diversity) and not from seeds (0.3-15%). The outcome of seed and soil microbiota coalescence is therefore strongly asymmetrical with a dominance of soil taxa. Interestingly, seedling microbiota was primarily composed of initially rare taxa (from seed, soil or unknown origin) and sub-dominant soil taxa. Our results suggest that plant microbiome engineering success based on native seed or soil microbiota will rely on rare and sub-dominant taxa in source communities.

## Introduction

Plants live in complex associations with a wide variety of microorganisms that can modulate their fitness (1–3). The plant microbiota is mainly acquired horizontally through different environmental sources including soil (4), air (5), rainfall (6) and insects (7). However, some members of the plant microbiota are also acquired vertically through vegetative propagation (8) or sexual reproduction *via* seeds (9).

In spermatophyte, seed-associated microbial community constitutes the primary inoculum for the next plant generation (10). The seed microbiota can have a crucial role for crop installation by modulating dormancy (11,12), germination (13,14), seedling development (15–17) and recruitment of plant symbionts (18). Seed transmission of some specific plant-beneficial or phytopathogenic microbial strains is well documented (19–23). However, little knowledge is available on the fraction of the plant microbiota that is acquired through seeds. Pioneer studies indicate that a fraction of seed microbiota persists during germination and emergence and actively colonizes seedlings (24–26). Still, most of these studies have been carried out in gnotobiotic conditions in absence of other environmental sources, like soil.

The encounter between two microbial communities and how they then evolve together has been called community coalescence (27–29). The outcomes of community coalescence can vary in a continuum between two categories from symmetric to asymmetric outcomes (30). In symmetrical outcomes, the initial communities contribute quite equally to the resultant coalesced community, whereas in asymmetrical outcomes one initial community becomes dominant if considering the prevalence of the members. In the latter case, the resident community is generally favored at the expense of the invasive community.

This resident advantage can be explained by (i) a higher population size of the resident community (i.e. mass effects) (31), (ii) prior use of resource and space (i.e. priority effects) (32) or (iii) local adaptation of the resident community to the reception niche (i.e. community monopolization effect) (33).

To date, seed or soil microbiota have been extensively studied in isolation but their coalescence, the assembly mechanisms at play remain broadly unexplored and their coalescence type remains to be characterized. Seed and soil coalescence present a singular situation if the seedling community is considered as the outcome community. The seed community is extremely reduced in terms of abundance compared to soil community, but its members have potentially a ‘home advantage’ as they have been selected by the plant. We therefore hypothesized that despite its lower microbial richness and diversity, seed microbiota being already present and more adapted to the plant environment, would benefit from priority effects over soil microbiota for the colonization of plant compartments (18). Nevertheless, adaptation to seed does probably not recover all features for adaptation to a growing seedling. The initial soil microbial diversity level is probably another key driver of seedling microbiota and persistence of seed-borne microbial taxa. Indeed, we propose that a soil of high microbial diversity would induce a weaker colonization of the seed microbiota on seedlings due to stronger competition and higher functional redundancy in this soil (34).

In the present work, we monitored the outcome of seed and soil communities’ coalescence during early seedling growth. More specifically, we investigated (*i*) what is the main source of microbial transmission to seedlings; (*ii*) does seed microbiota composition and soil diversity impact the assembly of seedling microbiota; and *(iii)* does the initial taxon abundance in the source influence its transmission success? To answer these questions, we selected seed lots of different genotypes of the winter oilseed rape *Brassica napus,* which harbored contrasting seed microbiota (35). We simulated a total of eight different coalescence events by sowing the four seed lots in two soils of different microbial diversities. Bacterial and fungal community structure was characterized by amplicon sequencing of the source seed and soil communities and of the roots and stems sampled at different seedling growing stages. This study gained new insights into the soil and plant drivers that influence the coalescence of soil and seed microbiota during the first steps of seedling microbiota assembly.

## Materials and Methods

### Soil preparation

In 2014, soil was sampled at a depth of 10-30 cm in a 20-year wheat experimental plot (La Gruche, Pacé, France, 48°08’24.5”N l°48’01.0”W). Soil was sterilized and recolonized with two levels of soil microbial diversity according to the experimental procedure described in Lachaise *et al.* (36). In short, a fraction of the soil sample was ground and sieved at 2 mm. Three sub-samples (900 g each) were soaked in 8L of sterile water. The resulting soil suspensions were 1:10 serially diluted in sterile water up to the 10^-6^. Another fraction (~80 kg) of the soil sample was ground, sieved at 4 mm and mixed with 1/3 washed sand. This soil was dispatched in 2.5 kg bags and gamma-irradiated (35 kGy, lonisos, France). Each bag was inoculated with 320 mL of undiluted or 10^-6^ or soil suspensions, therefore resulting in soil with high and low microbial diversity, respectively. The soil recolonization was repeated 3 times (A, B, C). The bags were mixed and aerated in sterile conditions twice a week during the 39 incubation days at 20°C, to homogenize gaseous exchanges and recolonization. Microbial recolonization dynamics of soil was monitored by sampling 30g of soil at several times. For bacteria, each sample was serial diluted and plated on 1/10 strength Tryptic Soy Agar(17g.L^-1^ tryptone, 3 g.L^-1^ soybean peptone, 2.5 g.L^-1^ glucose, 5 g.L^-1^ NaCI, 5 g.L^-1^ K_2_HPO_4_ and 15 g.L^-1^ agar) supplemented with Nystatin (0.025 g.L^-1^), and the number of colony-forming units was measured after 3 days of incubation at 27°C. For fungi, following the pourplate method, serial diluted samples were mixed with molten Acid Malt Agar (10 g.L^-1^ malt extract, 15 g.L^-1^ agar and 0.25 g.L^-1^ citric acid) supplemented with streptomycin (0.15 g.L^-1^) and penicillin (0.075 g.L^-1^) and the number of colony-forming units was measured after 7 days of incubation at 20°C. After 39 days of incubation, a portion of each soil (high and low diversity) in three independent repetitions (A, B, C) was collected and stored at −80°C until DNA extraction. Organic and mineral composition and pH of these samples were analyzed (INRAE, Arras, France). Differences between soil composition were considered as significant with T-test at a p-value <0.1.

### Sowing and sampling

After 39 days of soil incubation, individual pots were filled with a 5 mm layer of sterile vermiculite and 80g of soil of high or low microbial diversity. Soils were saturated with tap water by sub-imbibition one day before sowing, in order to reach approximately 80% of humidity (retention capacity) on the day of sowing. During the experiment, plants were watered twice (at 5 and 12 days) with tap water.

Seed samples from two genotypes of *Brassica napus* (Boston and Major) collected during two consecutive years (Y1 and Y2) were selected for this work, resulting in four contrasted initial seed microbiota used for the coalescence experiment. At d0, seeds of each genotype and year were individually sown at a depth of 5mm and grown under controlled conditions (l4h day/ 10h night period, 20°C). Seven days (d07) and fourteen days (d14) after sowing, 10 and 20 plants were sampled per modality (2 genotypes x 2 harvesting years x 2 soils), respectively. Roots were cut from stems and gently soaked 10 seconds in sterile water to remove residual soil. Otherwise, stems were size-equalized on 2cm from the root basis. Therefore, the resulting sampled root habitat was composed of the inner root and rhizoplane and the stem habitat was composed of inner stem and stem surface. The 10 (d07) or 20 (d14) roots and stems were pooled separately and stored as root and stem compartments at −80°C until DNA extraction.

### Microbial DNA sample preparation

Seed samples were prepared for extraction as previously detailed in Rochefort et al., (2019). Briefly, seeds were soaked in phosphate-buffered saline (PBS, Sigma-Aldrich) supplemented with Tween^®^ 20 (0.05 % v/v, Sigma-Aldrich) during 2h30 at 4°C. DNA extraction was performed with the DNeasy PowerSoil HTP 96 Kit (Qiagen) following manufacturers procedure. Soil, root and stem samples were lyophilized before DNA extraction. Samples were mixed with 1mm and 3mm beads, and crushed 2 x 30s at 5m/s with FastPrep (MP Biomedicals). DNA extraction was performed with the DNeasy PowerSoil HTP 96 Kit.

### Library construction and sequencing

Amplicon libraries were constructed with the primer sets gyrB_aF64/gyrB_aR553 (24) and ITS1F/ITS2 (37), following the experimental procedure previously described in Rochefort *et al.* (35). A blank extraction kit control and a PCR-negative control were included in each PCR plate. Amplicon libraries were mixed with 10% PhiX and sequenced with two MiSeq reagent kit v3 600 cycles.

### Sequence processing

Primer sequences were removed with cutadapt version 1.8 (38). Fastq files were processed with DADA2 version 1.6.0 (39), using the following parameters: truncLen=c(190, 200), maxN=0, maxEE=c(1,1), truncQ=5 for *gyrB* reads. ITS1 reads were processed with the same parameters, except that reads were not trimmed to a fixed length and that trunQ was set to 2. Chimeric sequences were identified and removed with the removeBimeraDenovo function of DADA2. Taxonomic affiliations of amplicon sequence variants (ASV) were performed with a naive Bayesian classifier (40) implemented in DADA2. ASVs derived from *gyrB* reads were classified with an in-house *gyrB* database (train_set_gyrB_v4.fa.gz) available upon request. ASVs derived from ITS1 reads were classified with the UNITE v7.1 fungal database (41).

### Microbial community analyses

Analyses of diversity were conducted with the R package Phyloseq version 1.26.0 (42). Since the primer set gyrB_aF64/gyrB_aR553 can sometimes co-amplify *parE,* a paralog of *gyrB,* the *gyrB* taxonomic training data also contained *parE* sequences. Hence, ASV affiliated to *parE* or unclassified at the phylum-level were removed. Sequences were aligned with DECIPHER 2.14.0 (43) and neighbor-joining phylogenetic trees were constructed with Phangorn 2.5.5 (44). Identification of sequence contaminant was assessed with decontam version 1.4 (45) using the “prevalence” method at a threshold of 0.1. Fungal ASVs unassigned at the phylum level were removed. For the ITS1 dataset, we created a hybrid-gene phylogenetic tree (i.e. ghost tree) (46) based the 18S rRNA gene sequence database (Silva 132) and ITS database (UNITE v8) in order to perform phylogenetic analyses.

Data were normalized based on sequencing depth. For soil analyses, data were rarefied at 30 000 and 25 000 reads per sample for *gyrB* and ITS1, respectively. For plant (root and stem) analyses, data were rarefied at 1 000 reads per sample for *gyrB* and ITS1 **(Fig. S1),** a validated choice to keep an optimal number of reliable samples. Faith’s phylogenetic diversity was calculated on *gyrB* and ITS1 datasets with the R package picante version 1.8 (47). Differences in alpha-diversity estimators were assessed with Analysis of Variance. Differences were considered as significant at a p-value < 0.05.

Changes in microbial community composition were assessed on log_10_+1 transformed values with Bray-Curtis (BC) index. Principal coordinate analysis (PCoA) was used for ordination of BC index. Homogeneity of group dispersions was assessed with betadisper function of vegan 2.5.3 (48). Difference in dispersion between groups was assessed with a Wilcoxon non-parametric test. To quantify the relative contribution of seed microbiota, soil diversity, development stage and habitat in microbial community profiles, permutational multivariate analysis of variance (PERMANOVA) (49) was performed with the function adonis2 of the R package vegan 2.5.3 (48). The relative abundance of ASVs belonging to ammonia-oxidizing bacteria (AOB) and nitric-oxidizing bacteria (NOB) according to their taxonomic classification were investigated in soils of high and low microbial diversities. These include *Nitrosospira* sp. and *Nitrosovibrio* sp. for AOB and *Nitrobacter sp.* and *Nitrospira sp.* for NOB (50,51). Differences in relative abundance of AOB and NOB were considered as significant with T-test at a p-value <0.1.

### Transmission analyses

A prevalence matrix of common ASVs between seeds, soils, roots and stems was constructed on non-rarefied data to have the most exhaustive presence-absence analysis of all taxa. Owing to the microbial variability between individual seeds, an ASV was recorded as present if detected in at least one of the 3 seed sample repetitions. Visualization of bacterial and fungal ASVs shared between biological habitats or specific to one was assessed with the R package UpSetR 1.4.0 (52).

The datasets supporting the conclusions of this article are available in the European Nucleotide Archive under the accession number PRJEB41004.

## Results

### Selected seed lots have distinct microbial communities

Harvesting year and plant genotypes are significant drivers of the diversity structure of the *B. napus* seed microbiota (35). Two *B. napus* genotypes (Boston and Major) collected during two harvest years (Y1 and Y2) were selected for this work. The most prevalent seed-borne bacterial ASVs were affiliated to *Sphingomonas* and *Frigoribacterium* genera **(Fig. S2A).** Fungal communities were dominated by ASVs affiliated to *Cladosporium* and *Alternaria* genera **(Fig. S2B).**

Estimated bacterial richness (Chao1) was on average at least two times higher in Y2 (250 ASVs) than in Y1 (100 ASVs) for both genotypes, and phylogenetic diversity (Faith’s PD) was also significantly (p < 0.05) higher in Y2 **(Fig. 1A).**

**Figure 1.**
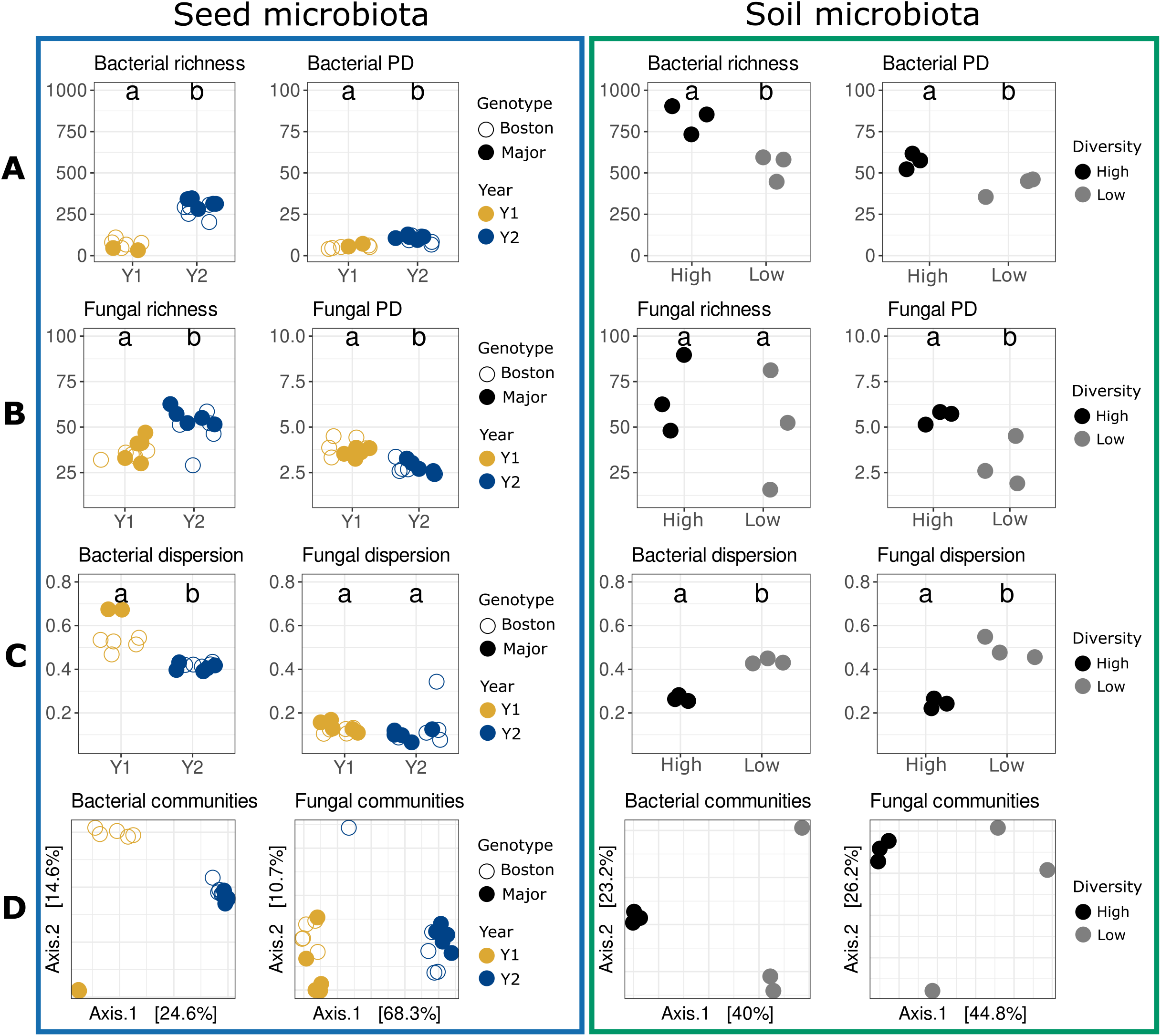
Diversity of initial seed and soil microbial community pools before coalescence. (A) Bacterial alpha-diversity. (B) Fungal alpha-diversity. (C) Microbial dispersion to centroid. (D) Microbial beta-diversity. Bacterial and fungal richness were estimated with Chao1. Bacterial and fungal phylogenetic diversities were calculated with Faith’s phylogenetic diversity (PD). PCoA ordination of Bray-Curtis index was calculated for bacteria and fungi. Seed microbiota (blue) was described in two *B. napus* genotypes (Boston, Major) harvested during two consecutive years (Y1, Y2). Soil microbiota (green) was described in two soils originating from dilution-extinction serials (High-, Low-diversity).

While estimated fungal richness was also higher in Y2 (~50 ASVs) than in Y1 (~35 ASVs), phylogenetic diversity was higher in Y1 **(Fig. 1B).** According to distance from centroid, variation in bacterial and fungal community composition (BC index) was significantly lower (p < 0.05) in Y2 in comparison to Y1 **(Fig. 1C).** Structure of microbial communities was significantly (p <.001) impacted by harvesting year with 24% and 62% of variance explained by this factor for bacteria and fungi, respectively **(Fig. 1D).** Moreover, seed genotype was also significantly (p < 0.001, 13% of variance) impacting bacterial community composition but not fungal community composition **(Fig. 1D).** In brief, the structure of microbial communities was different between the seed lots selected for this study and harvest year was the most important driver of these changes.

### Production of soils with contrasting levels of diversity

To obtain two soils with contrasted levels of microbial diversity, gamma-irradiated soil was inoculated with undiluted and diluted soil suspensions (Materials & Methods). After 39 days of incubation, a plateau of 10^9^ bacterial CFU and 10^5^ fungal CFU per gram of soil was reached for each soil **(Fig. S3).** The acidity level of low-diversity soils significantly (p = 0.003) increased by 0.65 pH unit compared to high-diversity soil. In addition the amount of nitric nitrogen was twice as large in high-diversity soils (p < 0.001), whereas ammoniacal nitrogen significantly (p < 0.001) decreased **(Table 1)**. Relative abundance of AOB was significantly (p < 0.1) higher in the soil of high diversity in comparison with the soil of low diversity. Moreover, no NOB was detected in the soil of low diversity while the soil of high diversity was composed on average of 0.25% NOB.

**Table 1:** Soil physicochemical analyses and relative abundance of bacterial nitrifiers. Values for high and low soil microbial diversities are the average of the 3 replicates ± standard errors. T-test was performed on data, and significant p-values are indicated with an asterisk. AOB = Ammonia-oxidizing

| Soil diversity | pH | Organic carbone (C) g/kg | Total nitrogen (N) g/kg | Organic material g/kg | C/N ratio | Total calcium carbonate (CaCO<sub>3</sub>) g/kg | Nitric nitrogen (N of NO<sub>3</sub>) on extract 1/10 mg/kg | Ammoniacal nitrogen (N of NH<sub>4</sub>) on extract 1/10 mg/kg | AOB (%) | NOB (%) |
| --- | --- | --- | --- | --- | --- | --- | --- | --- | --- | --- |
| **High** | 6.24<br>± 0.03 | 7.30<br>± 0.32 | 0.84<br>± 0.04 | 12.6<br>± 0.56 | 8.65<br>± 0.20 | <1 | 72.2<br>± 0.44 | 1.96<br>± 0.42 | 5.7<br>± 2.7 | 0.25<br>± 0.03 |
| **Low** | 6.89<br>± 0.09 | 7.26<br>± 0.17 | 0.84<br>± 0.02 | 12.6<br>± 0.31 | 8.61<br>± 0.05 | <1 | 29.03<br>± 1.68 | 32.57<br>± 0.93 | 0.5<br>± 0.11 | 0 |
| **T-test** | 0.003* | 0.87 | 1 | 0.93 | 0.73 | 1 | 0.0002 * | 2.97.10<sup>-5</sup> * | 0.08* | 0.005* |

bacteria. NOB = Nitrite-oxidizing bacteria.

The most prevalent soil-borne bacterial ASVs were affiliated to *Massilia, Nitrosospira* and *Sphingomonas* **(Fig. S2A).** The most prevalent soil-borne fungal ASVs were affiliated to *Mortierella, Trichoderma* and *Exophiala* **(Fig. S2B).**

According to alpha-diversity indexes (Chao1 and Faith’s index), high-diversity soils presented a higher (p <0.05) bacterial richness (~800 ASVs) and phylogenetic diversity compared to low-diversity soils (~500 ASVs) **(Fig. 1A).** With regard to fungal communities, only phylogenetic diversity was greater in high-diversity soils **(Fig. 1B).** Variation in microbial community composition was more important for low-diversity soils compared to high-diversity soils **(Fig. 1C).** The initial level of dilution employed explained 40% and 39% of the variance observed in bacterial and fungal communities, respectively **(Fig. 1D).** Hence, the dilution approach employed in this work resulted in two soils with contrasted levels of phylogenetic diversity and distinct community structure.

### Initial level of soil diversity impacted the diversity and structure of the plant microbiota

The outcome of community coalescence between seed and soil microbiota was investigated on two plant compartments (stem and root) collected at two distinct plant developmental stages (d07 and d14). On the seedling roots and stems, the most prevalent bacterial and fungal ASVs presented abundance patterns clearly influenced by initial soil diversity (e.g. *Devosia* or *Fusicolla aquaeductuum,* **Fig. S2).** Diverse bacterial (e.g. *Afipia, Ensifer, Fictibacillus, Nocardiodes)* and fungal genera (e.g. *Peziza, Fusicolla, Metarhizium)* were dominant on seedling roots. The stems were dominated by bacterial ASVs affiliated to *Bacillus, Nocardioides, Devosia* and *Fictibacillus,* while the most prevalent fungal genera were *Fusarium, Fusicolla* and *Acremonium* **(Fig. S2).**

Seed genotypes and harvest years did neither impact estimated richness nor phylogenetic diversity in stem and root **(Fig. 2).**

**Figure 2.**
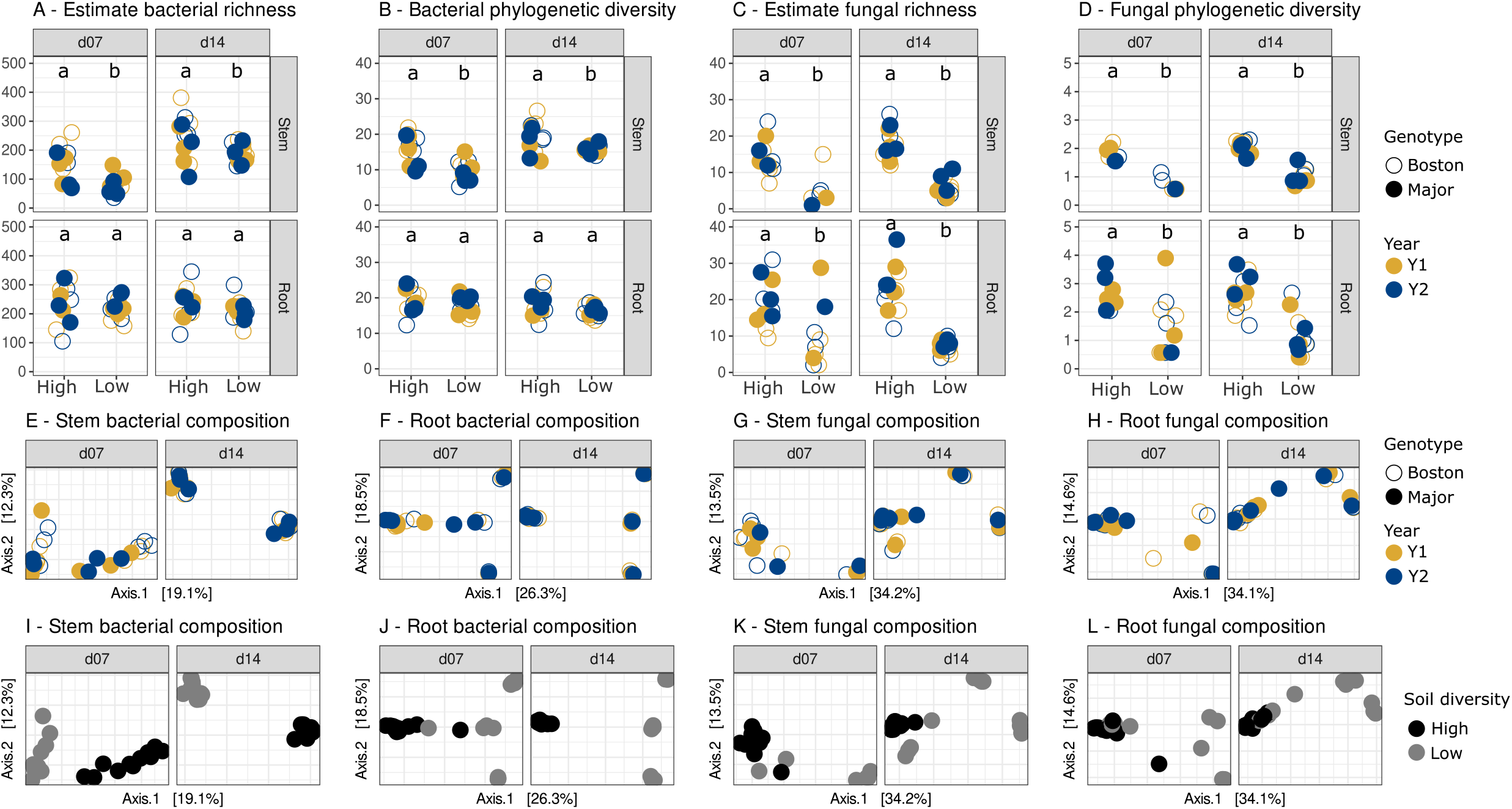
Structure of seedling microbiota after coalescence with contrasted seed and soil microbiota. (A, B) Bacterial alpha-diversity. (C, D) Fungal alpha diversity. (E, F, I, J) Bacterial beta-diversity in stems and roots. (G, H, K, L) Fungal beta-diversity in stems and roots. Open and closed dots are *B. napus* genotypes. Dots are colored either by harvest year (A-H) or by soil diversity (l-L). Bacterial and fungal richness were estimated with Chao1. Bacterial and fungal phylogenetic diversities were calculated with Faith’s phylogenetic diversity. PCoA ordination of Bray-Curtis index was calculated for bacteria and fungi.

In contrast, the initial level of soil diversity significantly (p <0.05) influenced microbial richness **(Fig. 2A** and **2C)** and phylogenetic diversity **(Fig. 2B** and **2D)** in stems at d07 and d14, with a higher richness and diversity in stems collected from high-diversity soils. No significant change in bacterial richness and phylogenetic diversity was detected in roots between the two soils **(Fig. 2A** and **2B)** while fungal richness and phylogenetic diversity were almost two times higher in roots from soil of high diversity at each development stage **(Fig. 2C** and 2D).

Moreover, there was a little but significant increase in bacterial richness and phylogenetic diversity in stems at stage d14 compared to stage d07. No differences were observed on fungal diversity across stages **(Fig. 2).**

Composition of the plant microbiota was not influenced by the seed genotype and the harvesting year **(Fig. 2E-H** and **Table 2).** In contrast, the initial soil diversity significantly (p < 0.001) contributed to the variance observed in stem (14.6%) and root (36.8%) bacterial communities **(Fig. 2I-J)** as well as the variance in stem (24.1%) and root (22.1%) fungal communities **(Fig. 2K-L** and **Table 2).** Stem- and root-associated microbial community compositions were also significantly impacted by the plant developmental stage **(Table 2).**

**Table 2:** Permutational multivariate analysis of variance on seedling roots and stems microbial beta-diversity. Linear model was built with the adonis2 function on data separated by root or stem compartment, integrating soil diversity (high and low), sampling stages (d07 and d14), GxY (interaction between plant genotype and year), and interaction between soil and GxY Significant values and their associated percentage of variance

|  |  | **Bacteria** |  | **Fungi** |  |
| --- | --- | --- | --- | --- | --- |
|  |  | P-value | % of explained variance | P-value | % of explained variance |
| **Root** | Soil | **0.001 \*** | **36.8** | **0.001 \*** | **22.1** |
|  | Stage | **0.001 \*** | **10.2** | **0.003 \*** | **8.0** |
|  | GxY | 0.955 | - | 0.999 | - |
|  | Soil:GxY | 0.990 | - | 0.996 | - |
| **Stem** | Soil | **0.001 \*** | **14.6** | **0.001 \*** | **24.1** |
|  | Stage | **0.001 \*** | **46.5** | **0.002 \*** | **8.6** |
|  | GxY | 0.394 | - | 0.986 | - |
|  | Soil:GxY | 0.695 | - | 0.878 | - |

(R<sup>2</sup>) are in bold and followed by an asterisk ( $p < 0.05$ ).

### Relative contribution of seed-borne and soil taxa to seedling microbiota

To describe the outcome of community coalescence between seed and soil microbiota during seedling growth, we characterized the proportion of bacterial and fungal ASVs from seed, soil or unknown origin that composed the stem and root seedling microbiota. A total of 39 bacterial ASVs and 16 fungal ASVs were shared between seeds, roots and stems **(Fig. 3).** Among these ASVs, 16 bacterial and 4 fungal ASVs were also found in soil samples.

**Figure 3.**
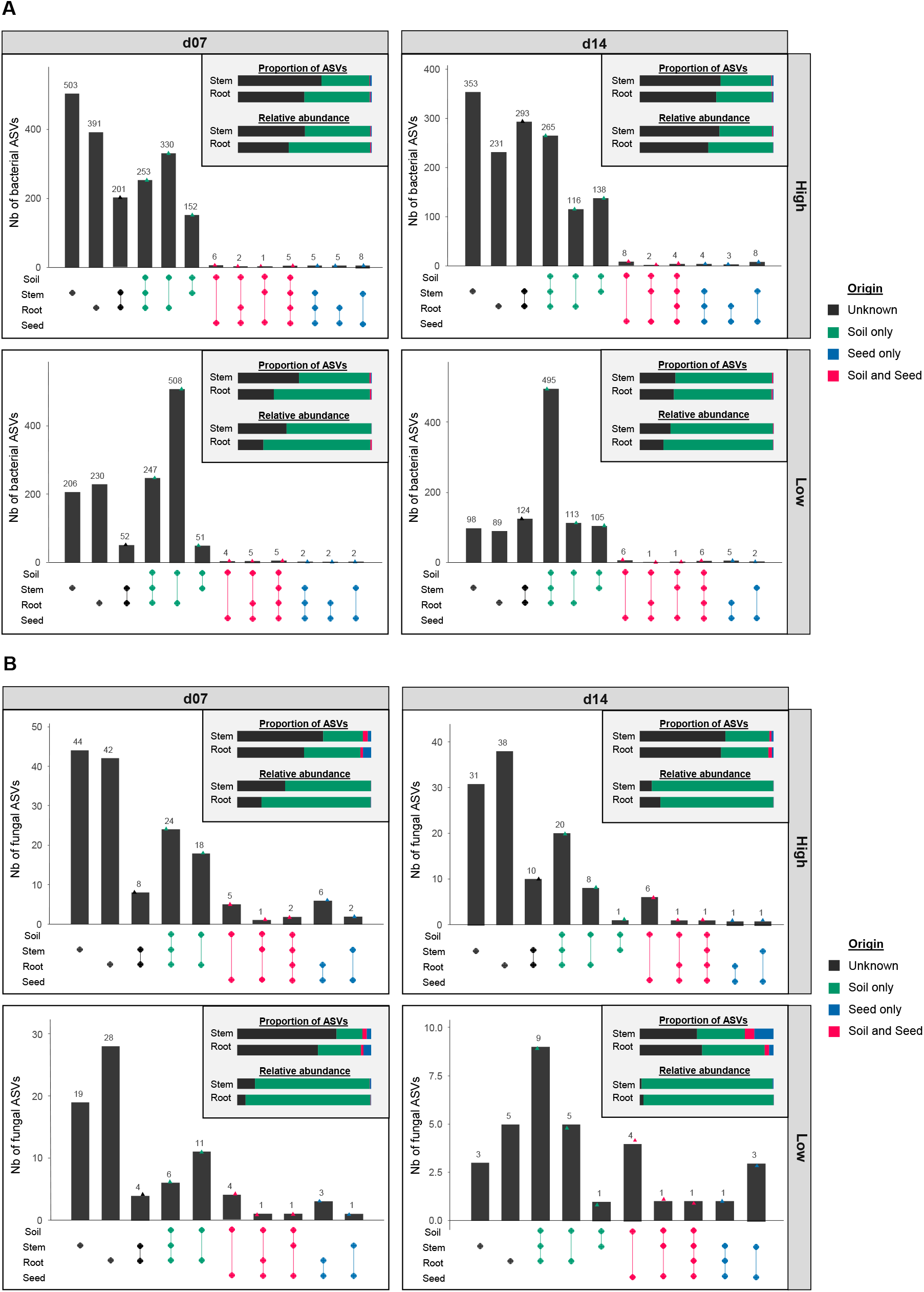
Origin of microbial taxa within stem and root assemblages. Bacterial (A) and fungal (B) ASVs shared between seeds, soil, roots and stems at each harvesting stage (d07 or d14) and soil diversity (High or Low). For each plot, upper-right bar charts summarize the proportion and relative abundance of plant-associated ASVs that were detected in soil and/or seeds or not detected (unknown) within these habitats.

Between 35 and 72% of seedling-associated bacterial ASVs were detected in the soil, while 20 to 45% of seedling fungal ASVs came from the soil **(Fig. 3).** These soil-derived ASVs represented between 40 and 98% of the total seedling microbiota. On the contrary, few seedling-associated ASVs were detected in the seed (0.3-15%) representing less than 1% of microbial relative abundance in roots and stems **(Fig. 3).** Lastly, a fraction of the seedling microbiota was of undetermined origin as undetected in the initial seed and soil pools. ASVs of undetermined origin either may correspond to unsampled taxa in soil or seed compartments (i.e. rare taxa) or may derive from other environmental sources. The proportion of bacterial ASVs of undetermined origin was higher for plants grown in the high-diversity soil (approximately 45-65% of community membership and composition) than in the low-diversity soil (20-30%) **(Fig. 3** and **Fig. S4A).** This result suggests that seedlings are primarily colonized by rare soil taxa that were initially undetected. In contrast, fungal ASVs not detected in soil and seeds represented approximately 40-60% of community membership in plants but their impacts on community *(i.e.* their relative abundance) was quite low with less than 20% regardless of the soil diversity **(Fig. 3** and **Fig. S4B).**

Contrasting microbial taxonomic composition was observed between the initial soil and seed pools. Concerning bacteria, Enterobacterales and Sphingomonadales were more abundant in seeds, while Burkholderiales were more abundant in soils **(Fig. 4A).** Changes in bacterial order relative abundance were also detected between stem and root, especially at d07 where Bacillales dominated the bacterial fraction of the stem microbiota **(Fig. 4A).** The taxonomic composition of the fungal fraction of the seed and soil microbiota was highly contrasted at the order level. Seeds were mainly inhabited by Pleosporales, Capnodiales and Tremellales, when soils were principally constituted of Mortierellales and Hypocreales **(Fig. 4B).** This latter order was however more prevalent in low-diversity soils in comparison to high-diversity soils **(Fig. 4B).** As in soil, Burkholderiales remained the dominant order in roots but were in part outcompeted by an increasing proportion of Rhizobiales. Xanthomonadales were reduced in roots especially in low-diversity soil and more drastically after 14 days. Bacillales were the transitory most dominant order at 7 days in stems before to regress at 14 days. On the opposite Pseudomonadales and Rhizobiales were transitory limited in stems especially in low-diversity soil before to increase at 14 days **(Fig. 4A).** For fungi, Hypocreales were highly prevalent in seedlings when Mortierellales were more or less prevalent at 7 days in roots and stems, then disappeared in roots and were not represented in stems at 14 days **(Fig. 4B).** These pools represented only a small proportion of transmitted ASVs from seed to seedling compared to the pools originating from soil **(Fig. 3).** Mainly bacterial ASVs belonging to Bacillales, Burkholderiales and Pseudomonadales and many fungal ASVs belonging to Hypocreales were transmitted from soil to seedlings.

**Figure 4.**
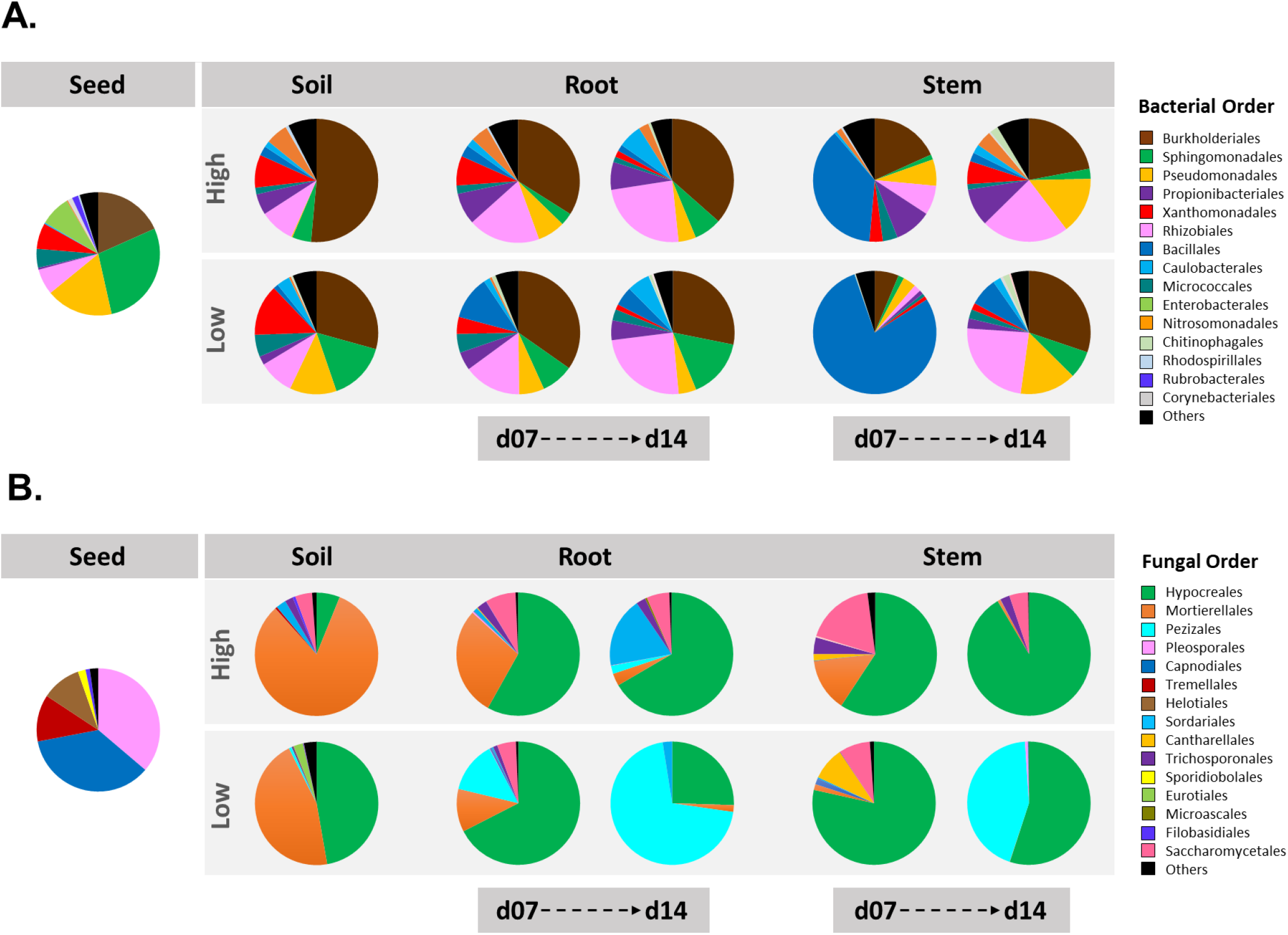
Taxonomic profiles of microbial communities associated with initial *B. napus* seeds and soil and with seedlings at 7 or 14 days after sowing. The 15 most abundant bacterial (A) and fungal (B) orders detected in seeds, soils, roots and stems are displayed.

### Seedling transmission success of seed-borne and soil microbiota

The contrasted seed lots and soils employed in the experimental design enabled the investigation in the influence of the initial diversity of the colonizing pools on the transmission success to the seedling **(Fig. 5).**

**Figure 5.**
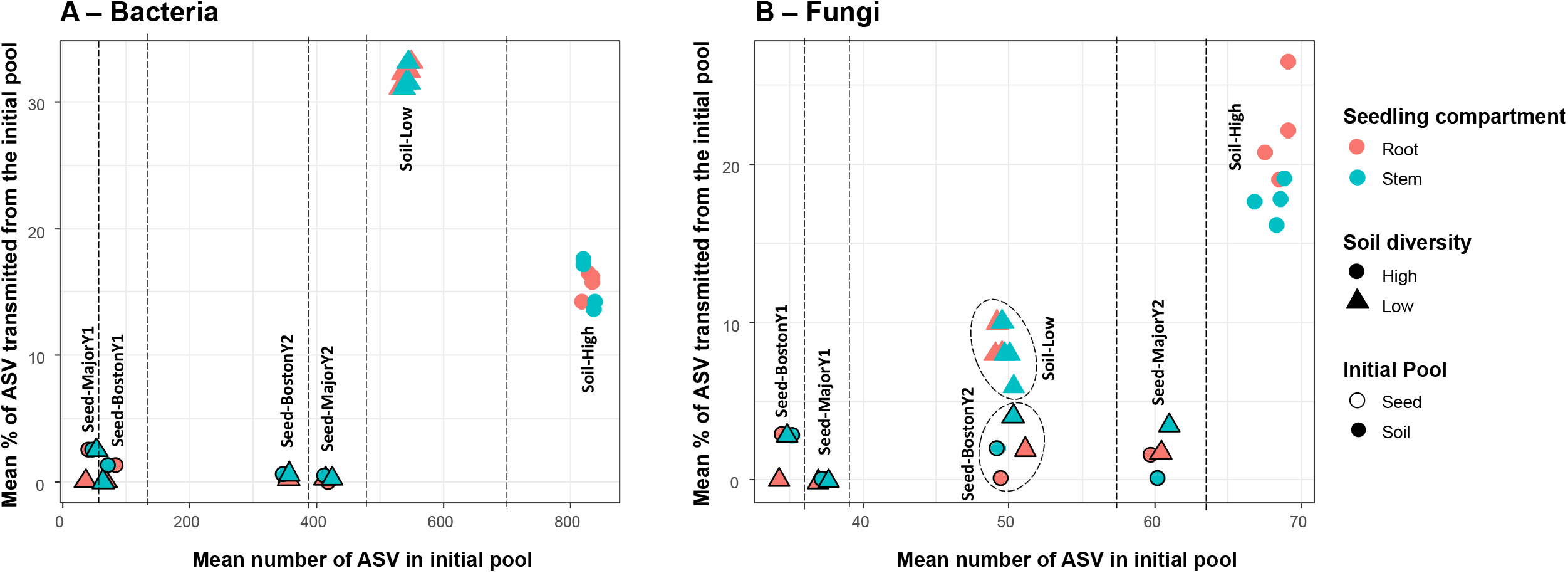
Influence of the initial bacterial (A) and fungal (B) diversity of the colonizing pools (seed or soil) on the transmission success to the seedling at d14. Nature of the initial pool (seed (Genotype/Year) or soil diversity (High-, Low-diversity)) is mentioned and separated with dashed lines. Colors represent the receiving seedling compartment (root or stem).

For bacteria and fungi, we found no clear relationships between the level of diversity of the initial pool (seed or soil) and the percentage of ASVs transmission in seedlings **(Fig. 5).** Soil microbiota always presented the highest rates of ASVs transmission to seedlings with 10 to 30% of the initial ASV pool. Overall, the transmission of soil-borne microbiota to seedlings was higher for the bacterial community (113-180 ASVs) than for the fungal community (3-18 ASVs, **Fig. 5A, 5B).** For bacteria, transmission to seedlings was higher in low-diversity soil (32% + 0.84) compared to high-diversity soil (15.6% + 1.49, **Fig. 5A).** For fungi, the opposite pattern was observed with a higher percentage of ASVs transmitted to seedlings from high-diversity soils (19.8% + 3.24) compared to low-diversity soil (8.5% + 1.41, **Fig. 5B).** A low percentage of ASVs transmission from seeds to seedlings was observed for bacteria (0.8% + 0.93) and fungi (1.45% + 1.44), with no clear influence of the genotype and harvest year **(Fig. 5).** Finally, overall ASV transmission to seedlings was similar on roots and stems, except for the fungal pool of the high soil diversity where transmission rate in roots was higher (22.1% + 3.18) than transmission rate in stems (17.7% + 1.2, **Fig. 5).**

### Emergence of rare and sub-dominant taxa in seedlings

*We* next examined the influence of the initial ASV relative abundance on its seedling transmission success and resulting abundance in roots and stems at d14. ASVs were grouped in three arbitrary abundance classes: rare (average relative abundance <0.01%), subdominant (between 0.01 and 1%) and dominant (>1%). An interesting finding was that dominant ASVs in the source pool (seed or soil) almost never became dominant in roots and stems (below 1:1 line) or were not even transmitted to seedlings **(Fig. 6).** The majority of transmitted seed-borne bacterial ASVs were initially rare taxa, with nine out 11 ASVs in roots and 11 out of 16 ASVs in stems **(Fig. 6A, 6C).** Only one dominant seed ASV (out of 65 in the initial pool) was found to be transmitted to roots and stems *(Afipia).* One sub-dominant ASV was successfully transmitted to both roots and stems *(Blastomonas)* and three sub-dominant ASVs to stems solely *(Pseudomonas lurida, Cutibacterium acnes, Hyphomicrobium)* **(Table SI).** It should be noted that none of these dominant and sub-dominant transmitted seed-borne bacterial ASVs became dominant in seedlings (relative abundance <0.3%).

**Figure 6.**
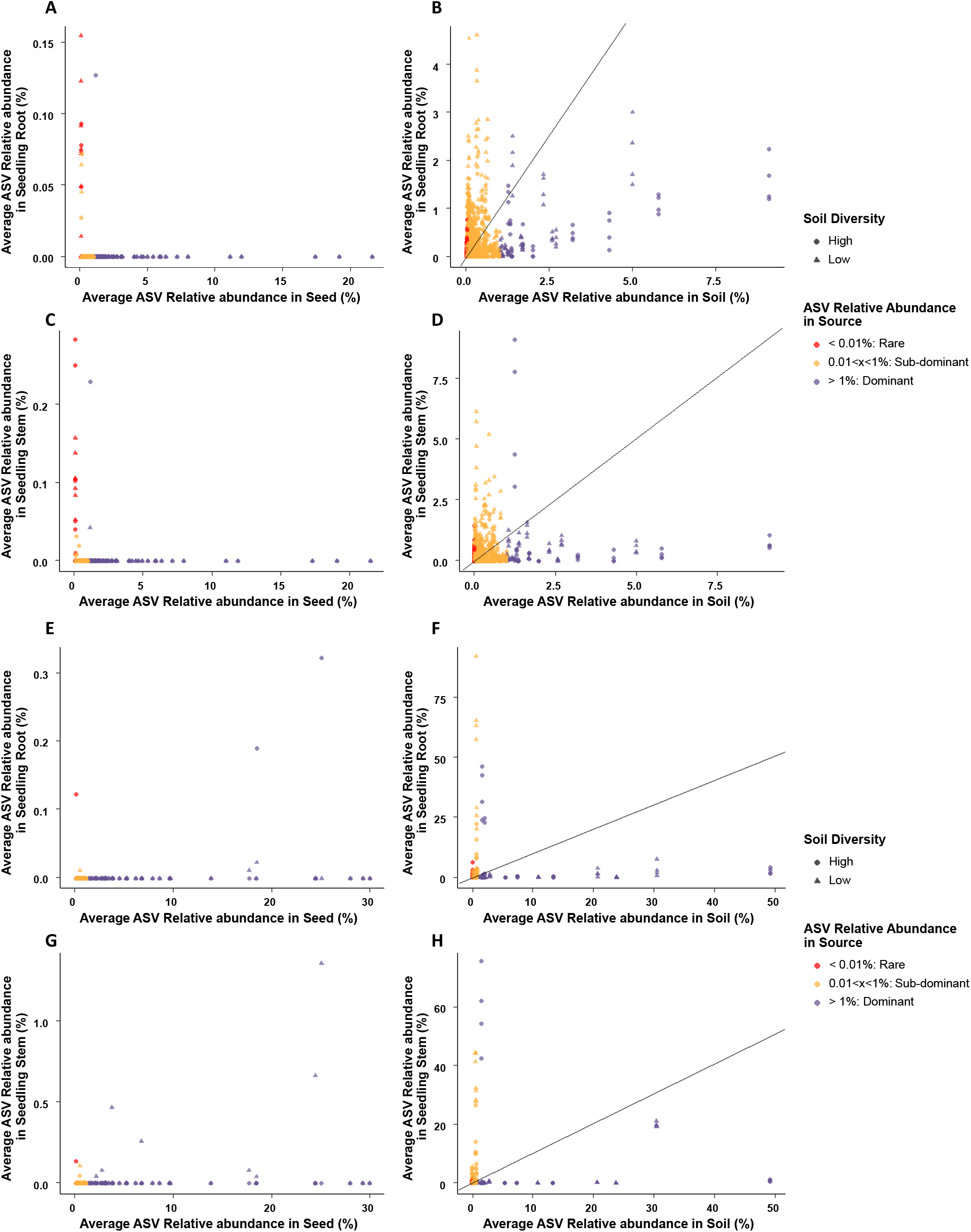
Relationship between relative abundance of transmitted bacterial (A-D) and fungal (E-H) ASVs in the source (seed or soil) and in seedling roots and stems. Each dot represents one ASV and is colored according to its relative abundance in the source (seed or soil).

The most abundant root bacterial ASVs (>1%) originated from soil-borne ASVs classified as sub-dominant ASVs (n=26 ASVs) and dominant ASVs (5 ASVs) **(Fig. 6B, 6D).** For the stems, most of the abundant ASVs originated from sub-dominant soil ASVs (n=24), a few dominant ASVs (n=6) and one rare ASV. When considering the entire seedling bacterial microbiota and not only the most abundant taxa, we found that 95% of the dominant soil ASVs, 40% of the sub-dominant ASVs and 28% of the rare ASVs were transmitted to roots. For the stems, 86% of dominant ASVs, 40% of sub-dominant ASVs and 29% of rare ASVs were transmitted.

For the fungal community, four seed-borne ASVs were transmitted to both roots and stems, composed of two dominant ASVs *(Alternaria infectoria, Cladosporium delicatulum),* one sub-dominant *(Alternaria sp.)* and one rare *(Gibberella avenacea).* In addition to these four ASVs, three additional dominant ASVs, all affiliated to A. *infectoria,* were transmitted to the stems **(Table S2).** For soil-borne fungal ASVs, the most transmitted taxa to roots were sub-dominant ASVs (19 ASVs, 12% of initial pool) with a few dominant taxa (9 ASVs, 53% of initial pool) and seven rare ASVs (12% of initial pool) **(Fig. 6E, 6F).** Similarly, soil-borne ASVs transmitted to stems were predominantly sub-dominant ASVs (15 ASVs, 10% of initial pool) along with seven dominant ASVs (41% of initial pool) and six rare ASVs (11% of initial pool) **(Fig. 6G, 6H).** It should be noted that some of these soil-borne fungal ASVs present extremely high relative abundance in seedlings (>20% relative abundance) compared to seed-borne ASVs (<2% relative abundance).

Altogether, these results indicate that seed-borne taxa transmitted to seedlings are predominantly rare taxa, while transmitted soil-borne taxa are primarily sub-dominant and dominant taxa. Moreover, this analysis shows that the transmission rates of fungi is two to four times lower than bacteria for all ASV abundance classes (dominant: 41-53% vs 86-95%; sub-dominant: 10-12% vs 40%; rare: 11-12% vs 28-29%).

## Discussion

### Asymmetric outcome of seed and soil community coalescence in favor of soil microbiota

Soil microbiota had a great influence on the plant microbiota composition since 70% and 30% of soil-borne bacterial and fungal taxa were detected in seedlings, respectively. In contrast, transmission of seed-borne microorganisms to roots and stems of *B. napus* was lower than soil with on average 2% of bacterial ASVs and 12% of fungal ASVs detected in seedlings. The remaining seedling-associated ASVs were not identified in both seed and soil habitats. It is highly plausible that most of these taxa were soil-borne and were initially not sampled due to the high heterogeneity of soil spatial structure (28). Indeed, there were on average many more bacterial ASVs of undetermined origin in the seedlings growing in the soil of high diversity (> 50%) than in those from a soil of low diversity (< 30%). These data suggest that a large part of these ASVs come from the soil but that they were not sampled in high-diversity soils. Overall, these results highlighted an asymmetric outcome of the coalescence between the soil and the seed microbiota (30), where the soil microbiota take precedence over the seed microbiota.

This asymmetric coalescence could be due the high population size of soil-borne taxa in comparison to seed-borne microbial population, a process known as mass-effect (31). Although seed microbial taxa can be considered as plant residents that are adapted to available niches (53), the successful invasion of root and stem by soil-borne taxa is undoubtedly facilitated by their population sizes. Alternatively, the dominance of soil associated microbial communities in seedlings could be explained by the higher amount of phylogenetic diversity in soil in comparison to seeds. Higher phylogenetic diversity could indeed increase the functional capabilities of soil associated microbial communities and then limit niche overlap between microbial entities (54). The level of diversity of the initial seed and soil pools were however not directly correlated to the number of seedling transmitted ASVs **(Fig. 5).** In other words, the diversity of the regional species pool (i.e. soils and seeds) were not a good predictor of the diversity of local communities (i.e. seedlings).

Weak transmission of seed-borne microorganisms to the rhizosphere has been previously reported in Asteraceae (55), Cucurbitaceae (56,57) and Poaceae (25,58). Survival on barley roots of seed-borne bacteria occurred in the non-growing part of the root system but not in emerging roots that are colonized principally by soil-borne bacteria (58). While we did not investigate the fate of seed-borne microorganisms in different parts of the root system of *B. napus,* the same proportion of seed-borne taxa were detected within stems or roots. Hence, the weak transmission of seed-borne taxa to seedlings was not a consequence of microhabitat heterogeneity.

### Seedling microbiota assembly is driven by the level of soil diversity but not the structure of the seed microbiota

The initial level of soil diversity was the key driver in the assembly of the stem and root microbiota. Indeed, stem microbial diversity and root fungal diversity were lower in the low-diversity soils compared to high-diversity soils **(Fig. 2).** Moreover, soil diversity explained between 15% to 37% of variance in microbial community composition in stem and root **(Fig. 2).** The impact of soil diversity on plant microbiota composition increased over time in roots and stems therefore confirming soil resident time and plant age in root microbiota dynamics (59,60).

The soil recolonization approach employed in this work did result in two soils with contrasted levels of bacterial and fungal diversity. This modification of soil microbial diversity also impacts the functional diversity and ultimately the soil physicochemical parameters. Changes in NH_4_^+^ and NO_3_^-^ concentrations and pH between low and high-diversity soils were indeed monitored. A similar drastic switch in the net balance between mineral nitrogen forms, from dominant nitrate forms in the high diversity soil to an altered N-cycle with dominant ammonium forms in the low diversity soil is reported in another soil dilution experiment (61). A lower NO_3_^-^ amount is consistent with a reduced abundance of bacterial and archaeal ammonia oxidizers. A higher NO_3_^-^ amount is linked with a decrease in the abundance of denitrifiers that can reduce NO_3_^-^ into N_2_. Nitrification process is associated with pH reduction prior to NO_2_^-^ accumulation period, with a potential decrease of 0.75 pH unit (62). Hence, the impact of soil diversity on seedling microbiota structure could not be solely due to the level of microbial diversity but to local changes of physicochemical parameters.

Contrary to what has been observed in the soil, we did not detect any significant impact of the initial seed microbiota composition on the overall structure of the stem and root microbiota. This observation reflected not only a low transmission of seed-borne taxa to roots and stems of *B. napus* but also a weak, if not an absence, of historical contingency. Historical contingency is mediated by priority effects that correspond to the impact of species on one another depending on its order of arrival within the local community (32). Importance of historical contingency in assembly of the plant microbiota was recently highlighted in wheat, where the identity of seed-transmitted fungal taxa can modify colonization of roots by dark septate endophytes, three weeks following germination (18). While we cannot conclude there is an absence of priority effects between microbial species within the root and stem microbial communities of *B. napus,* historical contingency is not promoted during seedling community assembly.

### Seedling microbiota is primarily composed of initially rare and sub-dominant taxa in source communities

The initial abundance of microbial taxa in the regional species pool was not positively correlated to their seedling transmission. This was particularly marked for bacterial ASV where rare seed-borne taxa and sub-dominant soil-borne taxa were mostly transmitted to the seedling **(Fig. 6, Fig. S5).** An increase in the abundance of rare taxa in seedlings could be explained by several attributes including more favorable environmental conditions for their growth and awakening from dormancy (63). Many seed-borne bacteria enter a viable but non-culturable (VBNC) state (64–66). Nutrient-rich environment, such as the spermosphere (67), can elicit the resuscitation of VBNC cells (68). In any case, the rare taxa that are selected on seedlings must carry specific genetic determinants that are responsible for their higher fitness that deserve to be investigated further. The emergence of rare taxa seems to be a shared coalescence outcome between different ecosystems, *i.e.* for the mixing of freshwater and marine microbiome (29), the unequal mixing of two soils with different physicochemical and microbial compositions (69) or the mixing of soils for the outcome of rhizobial communities on rooibos nodules (70). Rare taxa becoming dominant can provide essential or new functions in nutrient cycling or plant growth, which replace or compensate for the function deficiency of abundant species (71).

Interestingly, dominant taxa in the source pool (seed or soil) almost never became dominant in roots and stems. None of the dominant seed taxa (e.g. *P. agglomerans)* is transmitted maybe because these taxa are well adapted to ripe seed and low osmotic seed environment and not to growing seedling in competition with soil microbiota. Conversely, the better transmission of dominant and sub-dominant soil-borne taxa likely combines soil microbiota mass effect and a plant selective process among soil bacteria that are more adapted to root and stem habitats.

### Conclusion

In conclusion, we highlighted that the outcome of seed and soil microbiota coalescence was strongly asymmetrical with a clear dominance of soil microbiota. The level of soil diversity was an important driver of the structure of the seedling microbiota. Our approach enabled us to quantify the relative contribution of seed-borne and soil-borne taxa to seedling microbiota assembly and provided an estimation of transmission rates of each source microbiota. Out of all soil taxa detected, only 8 to 32% were able to colonize seedlings, while the proportion was of 0.8 to 1.4% for seed-borne taxa, which indicates a strong selection during seedling microbiota assembly. We demonstrate a high transmission of rare seed-borne and sub-dominant soil-borne taxa to seedlings, which is also a key feature of this coalescence **(Fig. S5).** These results provide an important foundation for the development of plant microbiome engineering through the modification of native seed or soil microbiota. Particularly, we encourage future work using natural or synthetic communities to integrate rare microbial taxa, as we showed that focusing only on dominant taxa in the source-communities is not informative to understand future seedling microbiota composition.

The community coalescence framework just started to emerge in the field of microbial ecology and we show here that this approach accelerates discoveries related to plant microbiota assembly and microbiome engineering. In this dynamic, further studies are needed to understand the ecological processes involved in this coalescence between seed and soil and how direct inoculation of microorganisms or modification of environmental conditions might alter its outcome.

## Supporting information

Supplemental informations

## Acknowledgements

We thank Muriel Bahut from the ANAN platform (SFR QuaSav) for amplicon sequencing and Laurent Philippot for discussion. This project named RhizoSeed was funded by INRAE (Metaomics and Ecosystems Metaprogram, Plant Health division) and Region Bretagne. We want to apologize to authors whose relevant work was not included in this article due to space constraints

## Code and data availability

R code and files for all microbiota analyses, statistics and figure generation are available on GitHub (https://github.com/arochefort/Seed_Soil_Coalescence).

