## Supplemental informations for "Asymmetric outcome of community coalescence of seed and soil microbiota during early seedling growth"

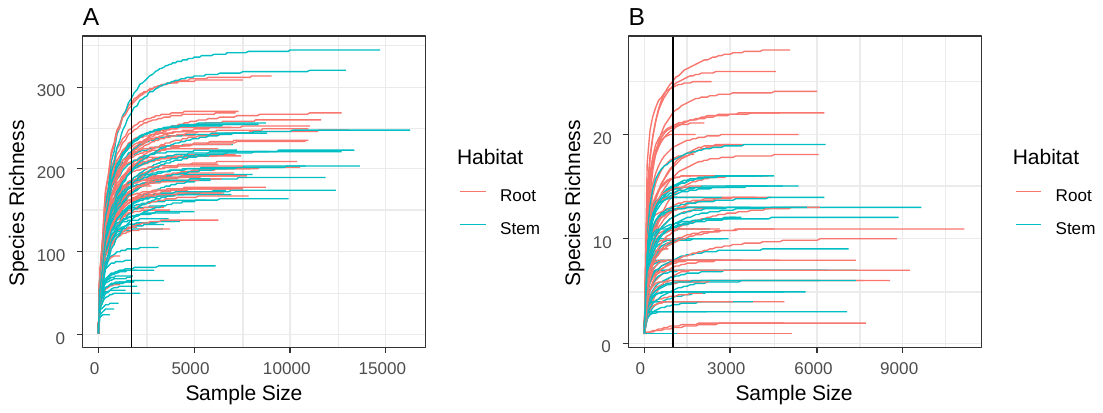

**Figure S1: Rarefaction curves of bacterial (A) and fungal (B) richness in root and stem samples.** Species richness represents the number of bacterial or fungal ASVs and sample size represents the number of reads. The vertical line represents the rarefaction threshold at 1000 read counts.

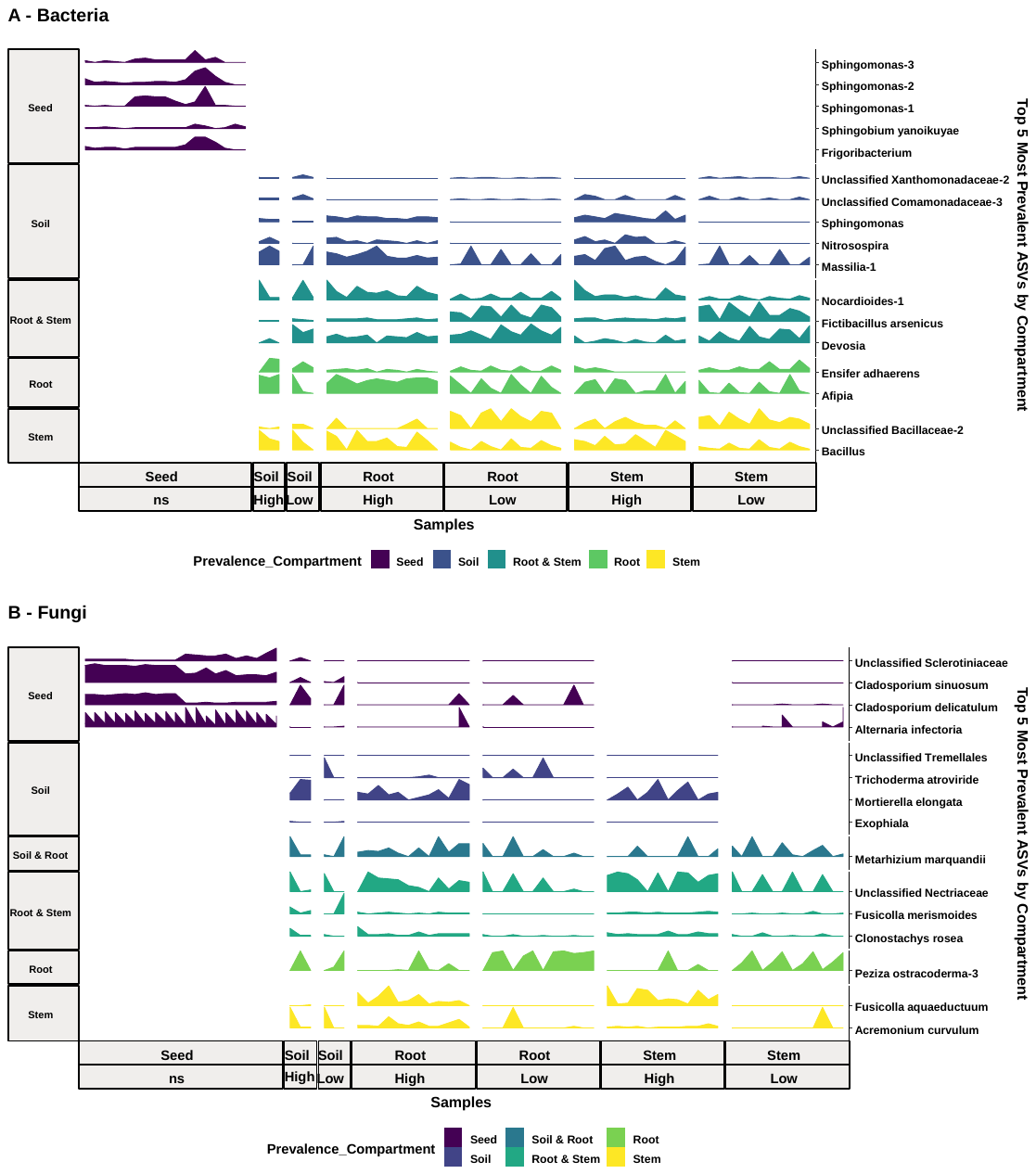
**Figure S2: Top 5 most prevalent bacterial (A) and fungal (B) ASVs by compartment and presence in the other compartments.**

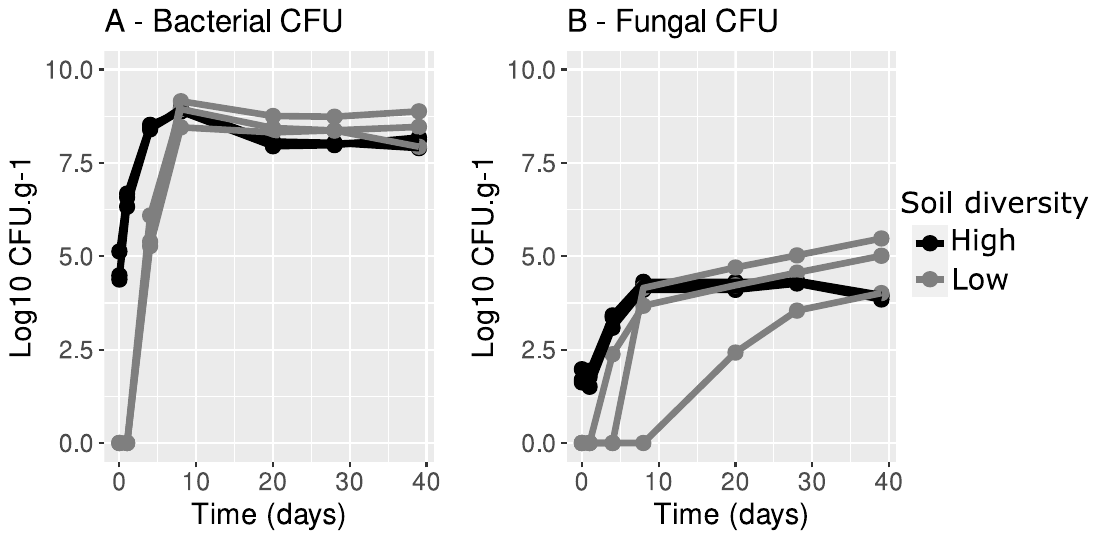

**Figure S3:** **Temporal soil recolonization by bacteria and fungi.** Bacterial and fungal populations are expressed as log10 CFU per gram of soil over time.

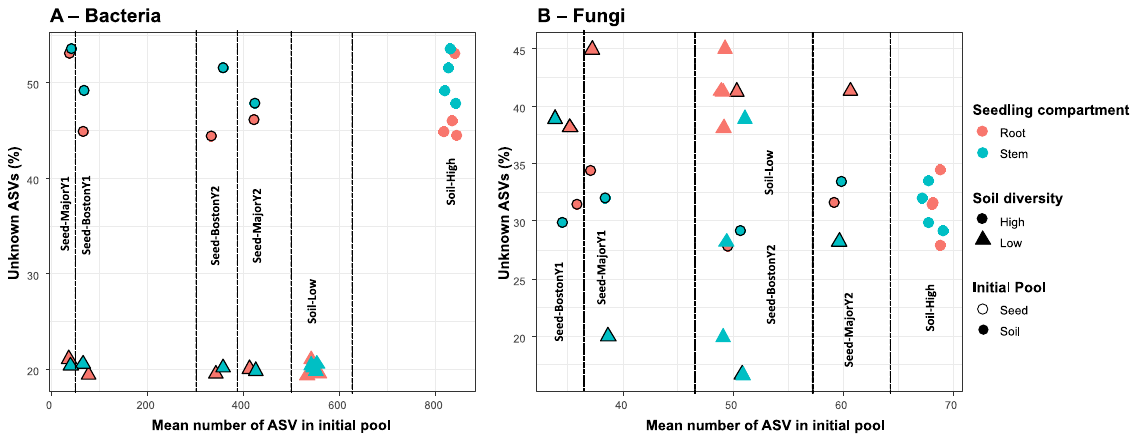

**Figure S4: Proportion of bacterial (A) and fungal (B) ASVs of unknown origin in seedlings according to diversity of the initial pool (seed or soil).** Nature of the initial pool (seed Genotype/Year or soil diversity) is mentioned and separated with dashed lines. Colors represent the receiving seedling compartment.

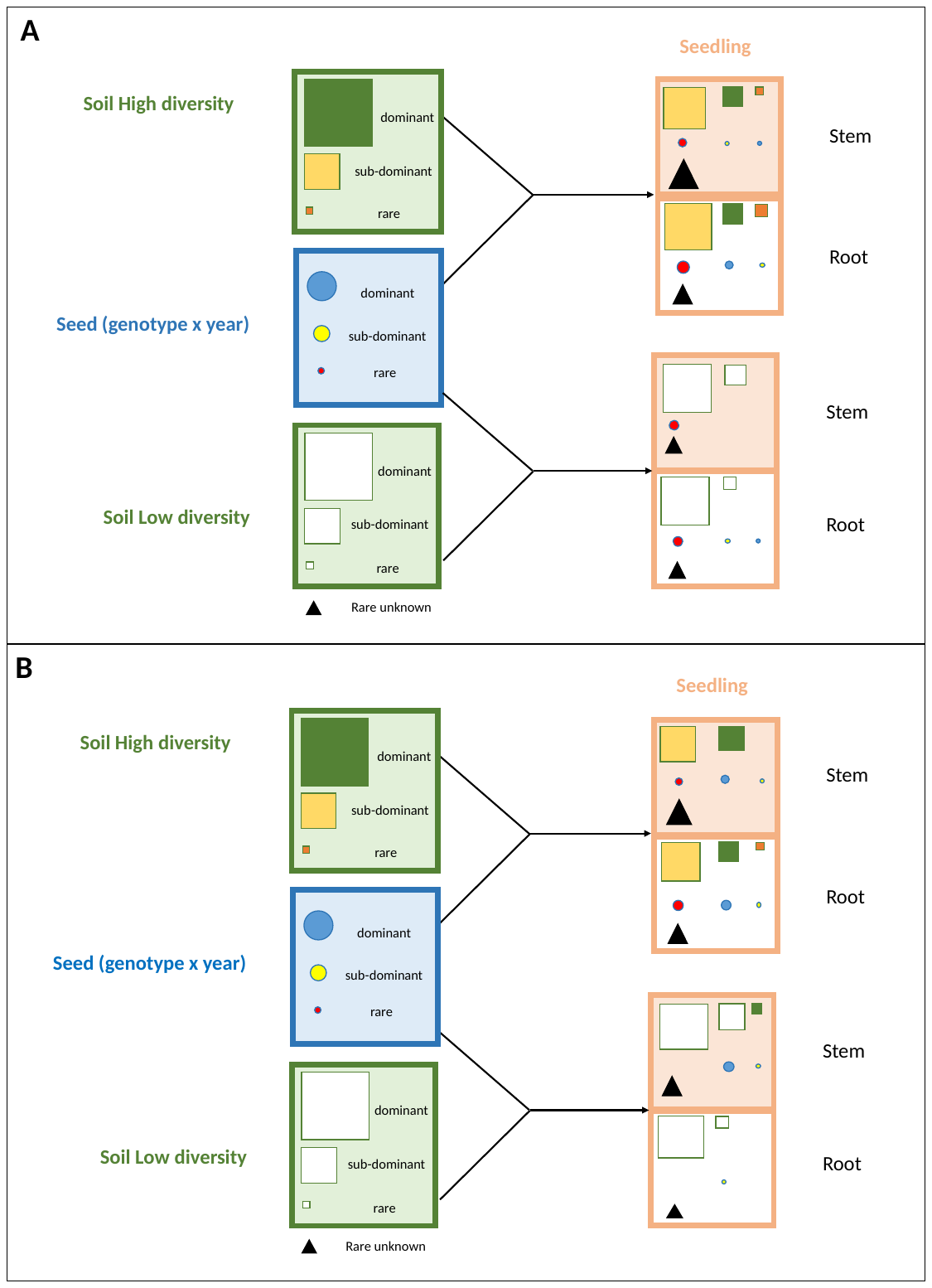

**Figure S5: Asymmetric outcome of coalescence of seed and soil microbiota during early seedling growth: relative abundance and transmission of bacterial (A) and fungal (B) taxa.**

**Table S1: Bacterial and fungal ASVs transmitted from seed to seedling (root and/or stem).** Relative abundance in seed: rare <0.01%, sub-dominant 0.01%<x<1%, dominant >1%.

| Bacteria |  |  |  |  |
| --- | --- | --- | --- | --- |
|  | **ASV** | **Order** | **Species** | **Relative abundance in seed** |
| root & stem | ASV314 | Micrococcales | *Arthrobacter sp.* | Rare |
|  | ASV448 | Pseudomonadales | *Pseudomonas sp.* | Rare |
|  | ASV458 | Micrococcales | *Terrabacter sp.* | Rare |
|  | ASV480 | Micrococcales | *Arthrobacter sp.* | Rare |
|  | ASV546 | Sphingomonadales | Erythrobacteraceae | Rare |
|  | ASV623 | Rhodobacterales | *Paracoccus yeei* | Rare |
|  | ASV757 | Sphingomonadales | *Blastomonas sp.* | Sub-dominant |
|  | ASV251 | Rhizobiales | *Afipia sp.* | Dominant |
| root only | ASV1000 | Micrococcales | *Arthrobacter sp.* | Rare |
|  | ASV1119 | Propionibacteriales | *Nocardioides sp.* | Rare |
|  | ASV4636 | Rhodobacterales | *Paracoccus sp.* | Rare |
| stem only | ASV273 | Bacillales | *Bacillus megaterium* | Rare |
|  | ASV334 | Xanthomonadales | *Stenotrophomonas rhizophila* | Rare |
|  | ASV419 | Burkholderiales | *Achromobacter sp.* | Rare |
|  | ASV651 | Micrococcales | *Arthrobacter sp.* | Rare |
|  | ASV2583 | Micrococcales | *Microbacterium sp.* | Rare |
|  | ASV479 | Propionibacteriales | *Cutibacterium acnes* | Sub-dominant |
|  | ASV1502 | Pseudomonadales | *Pseudomonas lurida* | Sub-dominant |
|  | ASV2520 | Rhizobiales | *Hyphomicrobium sp.* | Sub-dominant |

| Fungi |  |  |  |  |
| --- | --- | --- | --- | --- |
|  | **ASV** | **Order** | **Species** | **Relative abundance in seed** |
| root & stem | ASV170 | Hypocreales | *Gibberella avenacea* | Rare |
|  | ASV42 | Pleosporales | *Alternaria sp.* | Sub-dominant |
|  | ASV3 | Pleosporales | *Alternaria infectoria* | Dominant |
|  | ASV7 | Capnodiales | *Cladosporium delicatulum* | Dominant |
| stem only | ASV11 | Pleosporales | *Alternaria infectoria* | Dominant |
|  | ASV20 | Pleosporales | *Alternaria infectoria* | Dominant |
|  | ASV22 | Pleosporales | *Alternaria infectoria* | Dominant |

**Table S2: Bacterial and fungal ASVs transmitted by the soil and becoming dominant (>1%) in seedling.** Relative abundance in soil: rare <0.01%, sub-dominant 0.01%<x<1%, dominant >1%.

| **Bacteria** |  |  |  |  |
| --- | --- | --- | --- | --- |
|  | **ASV** | **Order** | **Species** | **Relative abundance in soil** |
| **root & stem** | ASV5 | Bacillales | *Fictibacillus arsenicus* | Sub-dominant |
|  | ASV8 | Burkholderiales | *Massilia sp.* | Sub-dominant |
|  | ASV13 | Pseudomonadales | *Pseudomonas sp.* | Sub-dominant |
|  | ASV20 | Rhizobiales | *Bosea sp.* | Sub-dominant |
|  | ASV22 | Rhizobiales | *Devosia sp.* | Sub-dominant |
|  | ASV30 | Propionibacteriales | *Nocardioides sp.* | Sub-dominant |
|  | ASV38 | Caulobacterales | *Caulobacter sp.* | Sub-dominant |
|  | ASV42 | Sphingomonadales | *Sphingopyxis sp.* | Sub-dominant |
|  | ASV49 | Sphingomonadales | *Sphingomonas sp.* | Sub-dominant |
|  | ASV51 | Rhizobiales | *Afipia sp.* | Sub-dominant |
|  | ASV54 | Burkholderiales | unclassified | Sub-dominant |
|  | ASV66 | Nitrosomonadales | *Methylobacillus sp.* | Sub-dominant |
|  | ASV4 | Burkholderiales | *Massilia sp.* | Dominant |
|  | ASV24 | Propionibacteriales | *Nocardioides sp.* | Dominant |
|  | ASV35 | Sphingomonadales | *Sphingomonas pruni* | Dominant |
| **root only** | ASV33 | Burkholderiales | Oxalobacteraceae | Sub-dominant |
|  | ASV36 | Rhizobiales | *Afipia sp.* | Sub-dominant |
|  | ASV43 | Burkholderiales | *Massilia sp.* | Sub-dominant |
|  | ASV68 | Sphingomonadales | Sphingomonadaceae | Sub-dominant |
|  | ASV71 | Burkholderiales | Comamonadaceae | Sub-dominant |
|  | ASV106 | Rhizobiales | *Bosea sp.* | Sub-dominant |
|  | ASV138 | Rhizobiales | *Pseudolabrys sp.* | Sub-dominant |
|  | ASV141 | Sphingomonadales | *Sphingopyxis sp.* | Sub-dominant |
|  | ASV145 | Propionibacteriales | *Aeromicrobium sp.* | Sub-dominant |
|  | ASV155 | Bacteroidetes | unclassified | Sub-dominant |
|  | ASV181 | Micrococcales | *Microbacterium sp.* | Sub-dominant |
|  | ASV227 | Burkholderiales | *Variovorax sp.* | Sub-dominant |
|  | ASV274 | Caulobacterales | *Phenylobacterium sp.* | Sub-dominant |
|  | ASV290 | Caulobacterales | *Caulobacter sp.* | Sub-dominant |
|  | ASV23 | Burkholderiales | *Massilia sp.* | Dominant |
|  | ASV39 | Sphingomonadales | *Sphingopyxis sp.* | Dominant |
| **stem only** | ASV359 | Alphaproteobacteria | unclassified | Rare |
|  | ASV12 | Pseudomonadales | *Pseudomonas fluorescens* | Sub-dominant |
|  | ASV14 | Pseudomonadales | *Pseudomonas sp.* | Sub-dominant |
|  | ASV26 | Burkholderiales | *Achromobacter sp.* | Sub-dominant |
|  | ASV37 | Burkholderiales | unclassified | Sub-dominant |
|  | ASV62 | Xanthomonadales | Xanthomonadaceae | Sub-dominant |
|  | ASV67 | Pseudomonadales | *Pseudomonas sp.* | Sub-dominant |
|  | ASV87 | Xanthomonadales | Xanthomonadaceae | Sub-dominant |
|  | ASV104 | Burkholderiales | unclassified | Sub-dominant |
|  | ASV128 | Pseudomonadales | *Pseudomonas moraviensis* | Sub-dominant |
|  | ASV137 | Rhizobiales | Hyphomicrobiaceae | Sub-dominant |
|  | ASV148 | Rhizobiales | *Devosia sp.* | Sub-dominant |
|  | ASV229 | Chitinophagales | Chitinophagaceae | Sub-dominant |
|  | ASV19 | Pseudomonadales | *Pseudomonas sp.* | Dominant |
|  | ASV34 | Pseudomonadales | *Pseudomonas fluorescens* | Dominant |
|  | ASV48 | Pseudomonadales | *Pseudomonas fluorescens* | Dominant |

| **Fungi** |  |  |  |  |
| --- | --- | --- | --- | --- |
|  | **ASV** | **Order** | **Species** | **Relative abundance in soil** |
| **root & stem** | ASV48 | Hypocreales | Nectriaceae | Rare |
|  | ASV55 | Hypocreales | *Dactylonectria novozelandica* | Rare |
|  | ASV63 | Hypocreales | *Neocosmospora rubicola* | Rare |
|  | ASV343 | Hypocreales | *Bionectria rossmaniae* | Rare |
|  | ASV477 | Diversisporales | *Entrophospora sp.* | Rare |
|  | ASV4 | Pezizales | *Peziza ostracoderma* | Sub-dominant |
|  | ASV7 | Capnodiales | *Cladosporium delicatulum* | Sub-dominant |
|  | ASV13 | Hypocreales | *Fusarium sp.* | Sub-dominant |
|  | ASV18 | Hypocreales | *Clonostachys rosea* | Sub-dominant |
|  | ASV29 | Hypocreales | *Fusicolla merismoides* | Sub-dominant |
|  | ASV34 | Trichosporonales | *Cutaneotrichosporon sp.* | Sub-dominant |
|  | ASV40 | Hypocreales | *Fusicolla aquaeductuum* | Sub-dominant |
|  | ASV47 | Hypocreales | *Acremonium curvulum* | Sub-dominant |
|  | ASV49 | Trichosporonales | *Trichosporon porosum* | Sub-dominant |
|  | ASV59 | Hypocreales | *Fusarium solani* | Sub-dominant |
|  | ASV62 | Hypocreales | *Volutella ciliata* | Sub-dominant |
|  | ASV96 | Hypocreales | *Gibberella intricans* | Sub-dominant |
|  | ASV141 | Hypocreales | *Metarhizium marquandii* | Sub-dominant |
|  | ASV170 | Hypocreales | *Gibberella avenacea* | Sub-dominant |
|  | ASV2 | Hypocreales | Nectriaceae | Dominant |
|  | ASV5 | Hypocreales | *Fusarium sp.* | Dominant |
|  | ASV6 | Mortierellales | *Mortierella elongata* | Dominant |
|  | ASV8 | Mortierellales | *Mortierella elongata* | Dominant |
|  | ASV30 | Sordariales | *Trichocladium asperum* | Dominant |
|  | ASV32 | Hypocreales | *Metarhizium marquandii* | Dominant |
|  | ASV35 | Trichosporonales | *Apiotrichum gracile* | Dominant |
| **root only** | ASV613 | Microascales | uncassified | Rare |
|  | ASV23 | Hypocreales | *Trichoderma atroviride* | Sub-dominant |
|  | ASV52 | Sordariales | *Chaetomium piluliferum* | Sub-dominant |
|  | ASV75 | Tremellales | uncassified | Sub-dominant |
|  | ASV146 | Hypocreales | *Clonostachys sp.* | Sub-dominant |
|  | ASV153 | Hypocreales | *Trichoderma velutinum* | Sub-dominant |
|  | ASV301 | Pezizales | *Peziza ostracoderma* | Sub-dominant |
|  | ASV17 | Mortierellales | *Mortierella alpina* | Dominant |
|  | ASV26 | Mortierellales | *Mortierella alpina* | Dominant |
| **stem only** | ASV11 | Pleosporales | *Alternaria infectoria* | Rare |
|  | ASV268 | Hypocreales | *Bionectria rossmaniae* | Sub-dominant |
